# Population density drives increased parasitism via greater exposure and reduced resource availability in wild hosts

**DOI:** 10.1101/2024.07.08.602460

**Authors:** Adam Z. Hasik, Shane Butt, Katie Maris, Sean Morris, Ali Morris, Richard S. Turner, Josephine M. Pemberton, Gregory F. Albery

## Abstract

Exposure to environmental parasites should increase with host population density due to the accumulation of infective parasites in space. However, competition for resources also increases with density, lowering condition and increasing susceptibility, which offers an alternative pathway for density-dependent infection to act. To test how these two processes act independently or together to drive greater parasite counts, we used a long-term study of red deer to examine associations between host density, resource availability, and counts of three common helminth parasites. We found that greater density correlated with reduced resource availability, and while density was positively associated with both strongyle and tissue worm burdens, resource availability was independently and negatively associated with the same burdens, supporting separate roles of density-dependent exposure and susceptibility in driving infection. This study provides evidence that competition for resources is an important driver of infection in higher-density areas, exacerbating the effects of density-dependent increases in exposure.

## Introduction

Parasites can impact organisms at every trophic level, affecting individual health, fitness, and interactions with other species (Ostfeld *et al*. 2018; De Lisle & Bolnick 2021; Acerini *et al*. 2022; Hasik & Siepielski 2022a; Hasik *et al*. 2023). As such, parasites can play an important role in regulating the size of host populations (Washburn *et al*. 1991; Hudson *et al*. 1998; Morand & Deter 2009). In addition to being widespread, parasites are unevenly distributed in space and time, such that host populations vary in both the prevalence (i.e., proportion of the host population parasitized) and intensity (i.e., mean number of parasites per infected host) of parasitism that they experience (Poulin *et al*. 2011; Gehman *et al*. 2017; Hasik & Siepielski 2022b). Such variation also occurs on the local scale, with fine-scale spatiotemporal variation in parasitism *within* host populations (Coltman *et al*. 1999; Forbes *et al*. 2012; Albery *et al*. 2022b). As a result, any parasite-mediated effects on host populations may also vary in space and time, which presents a challenge for quantifying if and how parasites regulate host populations. An important first goal to understanding the potential for parasite-mediated regulation of host populations is therefore to identify which variables best explain spatial variation in the distribution of parasites at fine spatial scales.

Host density is a central driver of fine-scale variation in infection; where more individuals inhabit a given area, they tend to encounter one another and their shed parasites more frequently, which can drive greater *per capita* rates of infection (Arneberg *et al*. 1998; Detwiler & Minchella 2009), a prediction central to understanding infectious disease dynamics (Côté & Poulin 1995; Altizer *et al*. 2003). However, parasite infection is a function of both exposure and susceptibility to infection, and population density can covary with environmental drivers that could influence these effects in either direction. For example, badger (*Meles meles*) population spatial structure limits exposure to areas of high parasite transmission, resulting in negative density-dependent infection rates (Albery *et al*. 2020a).

While density-dependent transmission has been the focus of disease ecology studies for decades, empirical evidence for density-dependent transmission is controversial and likely host-parasite system-specific (Brunner *et al*. 2017; Hopkins *et al*. 2020; Tompros *et al*. 2022), highlighting the need 1) to consider density-infection relationships in multiple host-parasite systems, and 2) to consider alternative mechanisms by which density can influence infection – one of which is the distribution of resources (Albery *et al*. 2020a; Albery *et al*. 2024).

Resource availability can covary with exposure and infection in important ways, determining the spatiotemporal distribution of infections. For example, because immunity is costly, increased nutrition and/or resource availability strongly improves host immune defenses (Boots 2011; Budischak *et al*. 2018; Hasik *et al*. 2021). In cases where immune function increases with resource availability (e.g., Boots 2011; Budischak *et al*. 2018; Hasik *et al*. 2021), parasite prevalence is predicted to decrease (Becker & Hall 2014). In this case, if high density correlates with high resource availability, it could counteract the density-dependent increases in exposure. However, if higher densities result in greater competition that outweighs the available resources, this could result in higher-density populations that are both more exposed and more susceptible, which could drive synergistic greater infection levels in these areas. Disentangling such environmentally-mediated processes is key to understanding parasite dynamics (Krasnov & Poulin 2010; Bolnick *et al*. 2020; Shearer & Ezenwa 2020; Hasik & Siepielski 2022b), and yet we have a very poor understanding of how resources, density, and infection covary and interact.

Density could covary with infection both positively and negatively in the wild depending on the balance of resources, competition, and foraging behavior. However, due to the scarcity of analyses of density-infection trends that take these processes into account with sufficient statistical power, it is unclear how these effects manifest. Specifically, density effects are often examined by comparing multiple populations – or a population at different times – to detect whether higher densities at the population level are associated with greater infection (McCallum *et al*. 2001; Lloyd-Smith *et al*. 2005; Hopkins *et al*. 2020). Recent work in Soay sheep showed that these global measures of density provide distinct information about the factors driving infection (Albery *et al*. 2024). In particular, correlated distributions of vegetation and host density in space and time are likely to drive divergent patterns of infection (Wiersma *et al*. 2023; Albery *et al*. 2024). Nevertheless, it remains unclear precisely how spatiotemporal variation in resource availability compares with the effects of density on parasite exposure.

Here, we use data from a long-term study of an exceptionally well-characterized wild ungulate population of red deer (*Cervus elaphus*) on the Isle of Rum, Scotland to ask how the local environment and host density best explain counts of three common parasite taxa. Although the population is known to be heavily spatially-structured both in terms of the distribution of parasites (Albery *et al*. 2019) and deer (Clutton-Brock *et al*. 1982; Albery *et al*. 2021b), we have yet to investigate how spatial structuring of parasitism might emerge from the distribution of population density, and how the effect of density might act via greater exposure compared to the effects of resource availability. We expected that higher individual parasite counts would be associated with greater density (through increased exposure), and, independently, with density-driven reductions in nutrition (through increased susceptibility). Untangling these two processes would help to identify a central cost of density for parasitism, expanding our understanding of density-dependence to encompass variation in resource availability.

## Material and Methods

### Study system

Data for this study were collected from a focal host population of red deer on the north block of the Isle of Rum, Scotland (57°N,6°20’W). Rum has a wet, mild climate and contains a mosaic of high-quality grassland and low-quality blanket bog and heath. The study area runs ∼4km north to south and ∼3km east to west with a total area of ∼12.7km^2^. The deer within the study area are wild, unmanaged, and free from predation pressure. The population is censused five times a month for eight months of the year along one of two alternating routes (Clutton-Brock *et al*. 1982), has a total of ∼250 individuals at any given time (Albery *et al*. 2018), and consists mostly of females and their recent offspring. Females have distinctive home ranges and live in loose social groups which often include matrilineal relatives (Clutton-Brock *et al*. 1982). The study area has heterogeneous vegetation, and the deer are mainly found on the grasslands bordering both sides of the Kilmory River (which runs north-south through the study area) and the coastal greens.

Since 2016, data on the helminth parasite burden of the population has been non-invasively collected by collecting fecal samples three times a year in April (Spring), August (Summer), and November (Autumn). In brief, observers note individually-recognized deer defecating from a distance and collect the fecal samples without disturbing the deer. Samples are then placed into plastic bags to keep the samples as anaerobic as possible and refrigerated at 4°C to prevent hatching or development of parasite propagules, with subsequent parasitological examination being conducted within three weeks in the case of strongyles (Albery *et al*. 2018). Detailed methods can be found in Albery et al. (2018).

Here we focus on three of the most common parasites infecting the red deer: strongyle nematodes (hereafter “strongyles”, a mix of different species with indistinguishable eggs), liver fluke (*Fasciola hepatica*), and tissue nematodes (*Elaphostrongylus cervi*). Strongyles have a direct lifecycle in which infective stages contaminate vegetation via fecal pellets and are subsequently consumed by a new host (Taylor *et al*. 2016). *F. hepatica* (Taylor *et al*. 2016) and *E. cervi* (Mason 1989) both have indirect lifecycles involving a snail intermediate host (the dwarf pond snail *Galba truncatula* and a number of land snails and slugs, respectively). After infecting and emerging from their intermediate hosts, larval *F. hepatica* contaminate vegetation near water bodies which are consumed by the deer final host. In contrast, deer become infected with *E. cervi* by consuming the intermediate snail host itself. While strongyle infections develop quickly such that calves excrete eggs within 2-3 months of birth, *F. hepatica* and *E. cervi* have longer prepatent periods, resulting in low prevalences of *F. hepatica* and *E. cervi* for juveniles relative to adults (Fig. S1).

### Measuring annual density

Census data were collected for the years 2016–2023, where individuals’ identities and locations (to the nearest 100m) were recorded. To understand the population-level correlates of parasitism in this system we focus on two individual-level biotic metrics. First, we calculated annual density using a previously described pipeline for this population (Albery *et al*. 2021b; Albery *et al*. 2022a) using all observations of each individual in each year. This approach uses a kernel density estimator, taking individuals’ annual centroids and fitting a two-dimensional smooth to the distribution of the data, producing a two-dimensional spatial distribution of the population. Individuals are then assigned a local density value based on their location on this kernel. Importantly, this is an individual-level measure of density as opposed to the usual population-level metric.

### Measuring resource availability

To investigate whether resource availability influences parasitism, we utilized Landsat satellite-derived measures of the Normalized Difference Vegetation Index (NDVI). NDVI values range from −1 to 1, with increasing positive values indicative of more biomass and/or greater productivity, making it a useful proxy of resource availability. NDVI is calculated as follows:

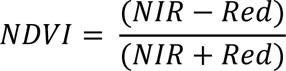

where ‘NIR’ and ‘Red’ are the respective wavelength values of the near-infrared and red spectral bands captured by the satellite’s optical sensor (Rouse *et al*. 1974; Pettorelli 2013). NDVI is widely-recognized as an accurate and reliable indicator of vegetation biomass and net primary productivity (Pettorelli *et al*. 2006; Pettorelli 2013), making it a suitable proxy of resource availability for herbivorous species.

We used the ‘LandsatTS’ package (Berner *et al*. 2023) to acquire Google Earth Engine-hosted Level-2 Collection-2 Tier-1 Landsat 5 (Thematic Mapper [TM]), Landsat 7 (Enhanced Thematic Mapper Plus [ETM+]), and Landsat 8 (Operational Land Imager and Thermal Infra-Red Scanner [OLI-TIRS]) satellite imagery. Throughout the study period, approximately one image of the study area was acquired per week, each with a pixel-level resolution of 30 meters. Each image was pre-processed to normalize surface reflectance and to categorize each pixel using the automated function of mask (fmask) algorithm (Zhu & Woodcock 2012). Prior to analysis, we excluded all pixels categorized as no data, cloud, cloud shadow, snow, or water, ensuring that only ‘clear’ ground surface pixels were used to calculate NDVI. We limited our analyses to pixels containing NDVI values of 0.15 and above, as values below this threshold typically indicate non-biomass areas such as concrete or buildings. Additionally, due to systematic differences in surface reflectance and spectral indices among Landsat sensors (Roy *et al*. 2016), we cross-calibrated the data among sensors to ensure that NDVI values were comparable regardless of the sensor. Cross-calibration followed the random forest model workflow of Berner et al. (2023, available from https://github.com/logan-berner/LandsatTS). In brief, the approach involved identifying the characteristic reflectance at sample sites during the growing season, defined as the beginning of March to the end of September, using Landsat 7 and Landsat 5/8 data from the same years. This was used this to train a random forest model to predict Landsat 7 reflectance from the Landsat 5/8 reflectance values. To account for a dearth of valid NDVI pixel data with which to train the model, we employed the high-latitude training dataset provided in the LandsatTS package. From these data, we again used a process from LandsatTS package by Berner et al. (2023) to quantify the growing season characteristics. This process involved iteratively fitting cubic splines to pixel measurements pooled over a seven-year moving window within the growing season. Observations were exponentially-weighted by distance in years from the focal year, so that observations from the focal year were most important in calculating its spline. Outliers were excluded and the splines refitted until all observations were within a 30% bound of the fitted spline. If there were fewer than ten observations in the focal window, the spline was not fit. We then estimated the maximum NDVI and the associated day of year for each pixel from its fitted spline. We assigned these values to the deer by matching the NDVI value of a given pixel to the mean annual centroid of each deer, giving us an estimate of the maximum amount of vegetation a given deer had access to during a given year. Because host density is not likely to be independent of resource availability, we tested for correlation between these two metrics (see *Results*).

### Do environmental and spatial factors explain parasitism?

To determine if host density and NDVI explained parasite counts among the red deer we used Integrated Nested Laplace Approximation (INLA) models. INLA models are a deterministic Bayesian approach which allow for the quantification of spatial effects and have been increasingly used for spatial analyses (Zuur *et al*. 2017; Albery *et al*. 2019; Albery *et al*. 2020a). We fit all models in R version 4.2.2 (R Core Team 2021) using the R-INLA package (Rue *et al*. 2009; Martins *et al*. 2013).

We constructed models for each of the three parasites using three different datasets. The first dataset included all deer for which we had parasite data (*n* = 4,272 records from *n* = 776 deer for the strongyles, *n* = 3,553 records from *n* = 729 deer for *F. hepatica*, and *n* = 3,559 records from *n* = 725 deer for *E. cervi*). The second dataset focused on juveniles up to 2 years old (*n* = 1,962 records from *n* = 617 deer for the strongyles, *n* = 1,622 records from *n* = 570 deer for *F. hepatica*, and *n* = 1,624 records from *n* = 567 deer for *E. cervi*). The third and final dataset included adult females only (*n* = 2,082 records from *n* = 242 deer for the strongyles, *n* = 1,703 records from *n* = 235 deer for *F. hepatica*, and *n* = 1,707 records from *n* = 234 deer for *E. cervi*). The lower sample sizes for *F. hepatica* and *E. cervi* are due to financial constraints in calendar year 2021 when they were not counted. In addition, no counts of any parasites were possible in spring 2020 due to pandemic restrictions (see Fig. S1 for a summary of parasite prevalence values for each season, year, and dataset). Investigating differences among all members of the population provided information on general trends across all age categories and sexes, while analyzing juveniles separately allowed us to understand if any patterns observed in all deer manifested early in life, where parasite-mediated effects on survival are apparent (Acerini *et al*. 2022). Adult females make up the majority of our data and focusing on them allowed investigation of how parasites relate to reproductive success.

For the models analyzing all deer we first fit a base model with age category (calf, yearling, two-year-old, and adult), season (spring, summer, and autumn), sex (male and female), and year (categorical) as covariates, while the models analyzing juveniles used the same base model structure except the age category variable only included calves, yearlings, and two-year-olds. For the models analyzing adult females only we fit a base model with season, age in years (a continuous variable), female reproductive status (none, meaning she had either never given birth or not in the year in question; summer, meaning she gave birth to a calf but it died during the summer; winter, meaning she gave birth to a calf that survived at least as far as the winter, and year (categorical) as covariates. In this system, female reproduction reduces immunity and increases parasitism (Albery *et al*. 2020b; Albery *et al*. 2021a), and increased parasitism reduces juvenile survival (Albery *et al*. 2021a; Acerini *et al*. 2022), in addition to adult survival and fecundity (Albery *et al*. 2021a), thus it is important to control for these differences in reproductive effort and age class. For each of our model sets, we then added to our base models by including annual density or NDVI alone or both terms together to understand if the inclusion of these variables would explain further variation in parasite counts.

Our focus in these analyses is on understanding if and how host density and resource availability explain parasitism in this population. Though each of the models described above for all deer and juveniles included important covariates such as age category, sex, and season due to their known relationships with parasitism in this system, we only present results on the relationships between annual density and NDVI and the abundance of each of the three parasites for the models analyzing these datasets. The full model results can be found in the Supplemental Materials for each of the three dataset.

## Results

Annual maximum NDVI was moderately negatively associated with annual density (R = - 0.33; Fig. S2). We also found that the counts of all three parasites varied in space across the study area. Fig. 1 shows the distributions of parasite count for all three parasites among all deer, as this was our largest and most inclusive dataset. In general, strongyle and *E. cervi* infections were most concentrated at the north end of the study area, yet heavier *F. hepatica* infections tended to be more concentrated toward the middle of the study area. Supporting the findings of previous studies in this system we found that parasite counts for all three taxa in the dataset containing all deer were highest in the spring and that both strongyle and *F. hepatica* counts were lowest in adults (Fig. S3). Strongyle and *F. hepatica* counts were similarly high in the adult females in the spring, but both *F. hepatica* and *E. cervi* counts moderately decreased as the females aged (Fig. S4).

**Fig. 1.**
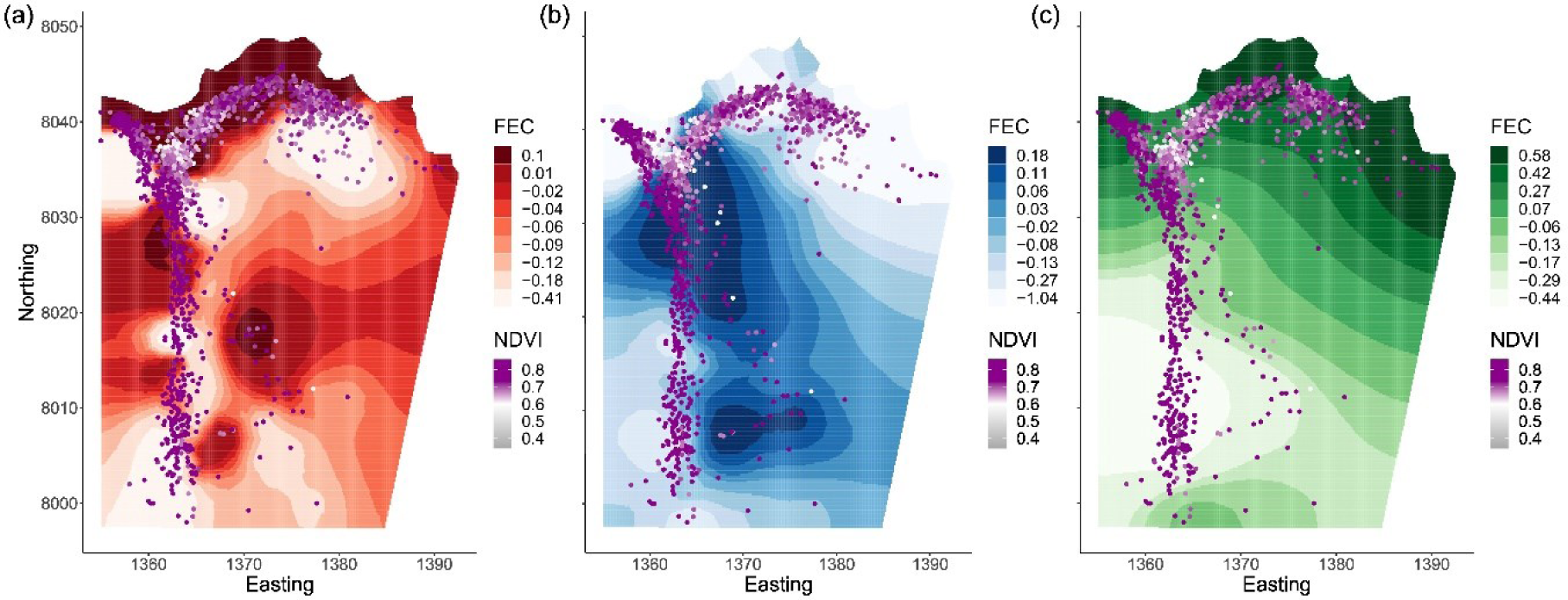
Spatial distribution of strongyle (a), *F. hepatica* (b), and *E. cervi* (c) counts throughout the study area, when considering data from all deer. Shown are projections of the spatially-distributed SPDE random effect from the null model for each parasite (providing a representation of where parasite counts are more abundant). Points represent the centroid of the estimated annual home range for a given deer, with color denoting the mean annual max NDVI a given deer was exposed to in a given year. Shading of the map denotes the lower bounds of nine quantiles of the spatial effects on the link scale, rounded to two decimal places, with darker colors representing higher parasite counts. Easting and Northing are in units of 100m grid squares, with 10 units equaling 1 km. The river at the base of the valley runs along the 1363 Easting.

Using the dataset that included all deer, we found that strongyle count slightly increased and decreased with annual density and NDVI, respectively (Fig. 2). These relationships were significant when the two factors were fit alone or together, though the strength of the relationship was reduced for both when they were fit together. Strongyle counts increased by ∼40% across the range of annual density (Fig. 3a), while strongyle counts were ∼2.4 times lower at the upper range of NDVI (Fig. 3b). NDVI was also negatively associated with *F. hepatica* counts, though only when it was fit together with annual density (Fig. 2b). In contrast to our findings for strongyles, we found a moderate negative association between *F. hepatica* counts and annual density (Fig. 2a), with the strength of the relationship increasing when annual density was fit together with NDVI. *F. hepatica* counts decreased by ∼66% and ∼78% across the range of annual density and NDVI, respectively (Fig. 3c-d). We did not find any significant relationships between the counts of *E. cervi* and annual density or NDVI, neither when they were fit alone nor together (Fig. 2).

**Fig. 2.**
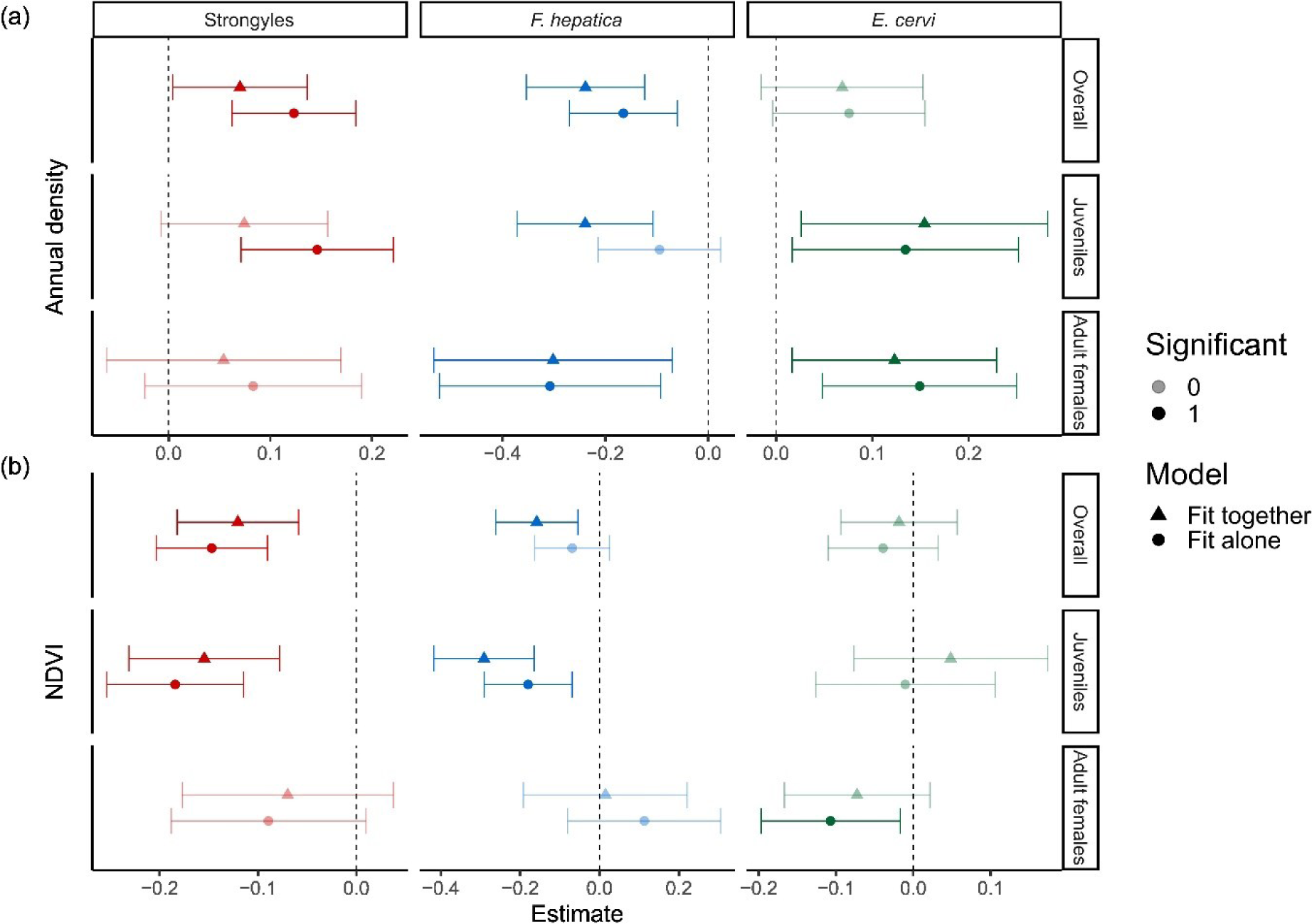
Forest plot of the relationships between individual annual density (a), individual mean annual max NDVI (b), and parasite counts for the datasets containing all deer, juveniles only, or adult females only, with panels for each parasite taxa and dataset. Points represent posterior estimates for mean effect sizes, error bars denote 95% credible intervals in standard deviations, color denotes the parasite taxa, and shape denotes whether an effect size comes from a model where the predictor was fit alone or together with the other main factor of interest. Significance of the effect size is denoted by the shading of the points.

**Fig. 3.**
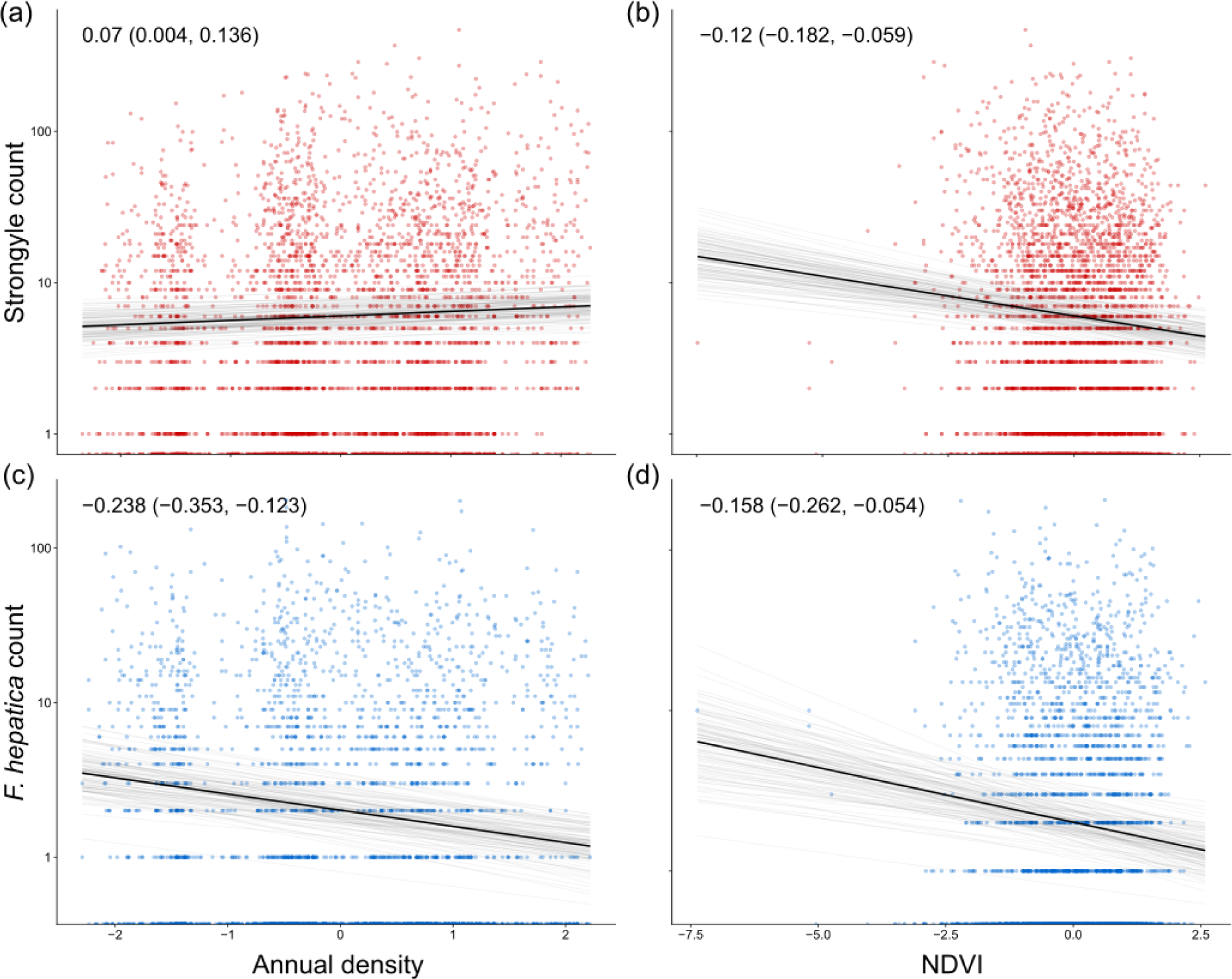
Strongyle and *F. hepatica* counts regressed on annual density (left columns) and NDVI (right column). The x axes denote individual annual density (a,c) and mean annual max NDVI (b,d), with parasite counts on the y axis on the log scale. All results are taken from the models fitting both predictors together for the overall dataset containing all deer, the dark black line represents the mean of the posterior distribution for the model estimates, the light grey lines are 100 random draws from the posterior to represent uncertainty. Points denote individual samples, with transparency to allow for visualization of overplotting.

The analysis of juvenile deer alone revealed similar patterns to those found in all deer (Fig. 2). However, for *F. hepatica* we found that the negative relationship with annual density was only significant when fit together with NDVI (Fig. 2a), while the relationship with NDVI was significantly negative when fit alone or together (Fig. 2b). We also found a weak positive relationship between annual density and *E. cervi* counts (fit alone or together with NDVI, Fig. 2a), though there was not a relationship with NDVI (Fig. 2b).

When considering adult females alone we found a moderate negative and weak positive relationship between annual density and *F. hepatica* and *E. cervi*, respectively (Fig. 2a), yet no relationship between density and strongyle count. NDVI was weakly negatively associated with the counts of *E. cervi* when NDVI was fit alone, yet we found no correlation with either strongyles or *F. hepatica* counts (Fig. 2b). *E. cervi* counts increased by ∼67% across the range of annual density (Fig. 4a), yet we did not find a significant relationship between *E., cervi* counts and NDVI (Fig. 4b).

**Fig. 4.**
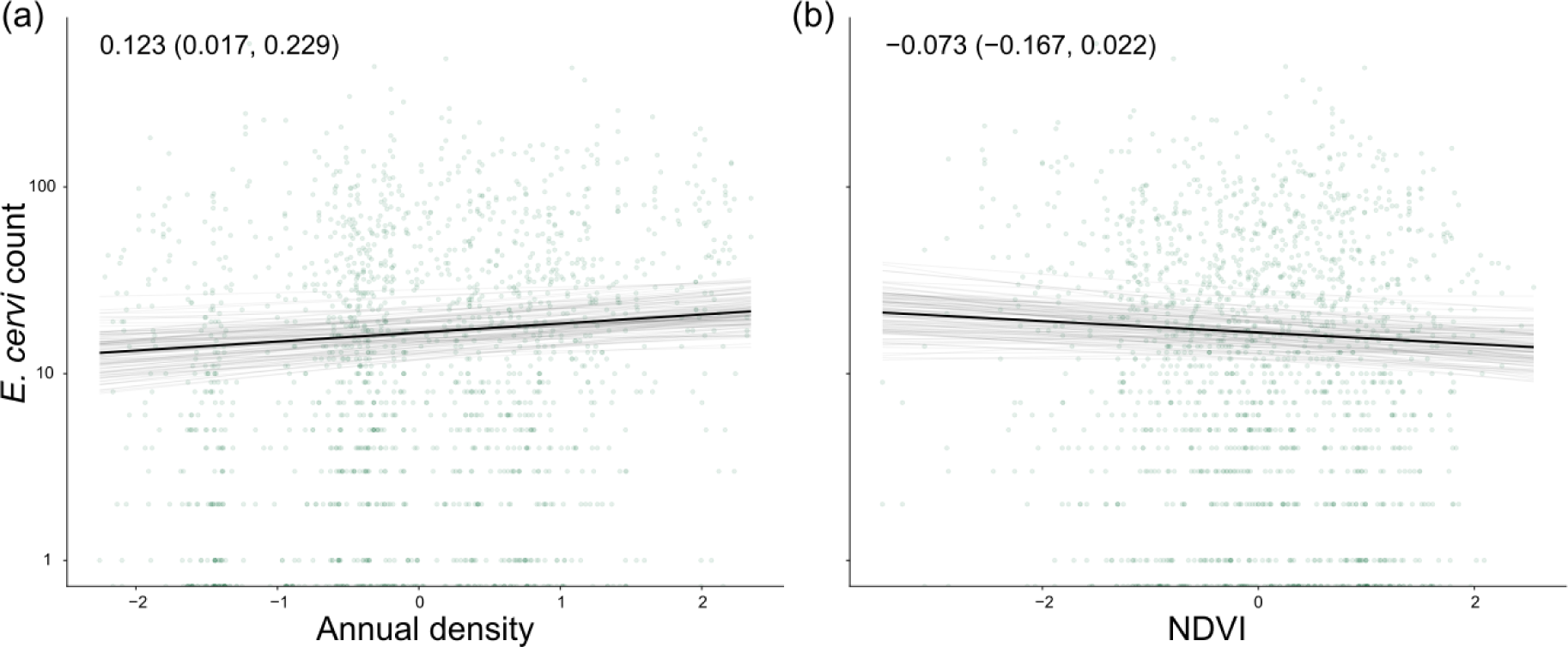
*E. cervi* counts regressed on annual density (a) and NDVI (b). The x axes denote individual annual density (a) and mean annual max NDVI (b), with parasite counts on the y axis on the log scale. All results are taken from the models fitting both predictors together for the dataset containing adult females only, the dark black line represents the mean of the posterior distribution for the model estimates, the light grey lines are 100 random draws from the posterior to represent uncertainty. Points denote individual samples, with transparency to allow for visualization of overplotting.

## Discussion

Understanding whether and how parasites regulate host populations remains an important goal, and a key first step is to link variation in the local environment and host ecology to host infection patterns. Here, we used data from an exceptionally well-characterized ungulate system to link host density and resource availability to infection with three different helminth parasite taxa. We found that deer experienced reduced resource availability at higher densities, and that higher density and lower resource availability were independently associated with increased counts of strongyles, with additional evidence for independent, positive associations between density and *E. cervi* counts in juveniles and adult females, and negative associations with *F. hepatica*. These findings highlight the twin roles of exposure and susceptibility in driving infection patterns (Sweeny & Albery 2022), and provide evidence that competition for resources and environmental confounding are likely important drivers of observed relationships between density and parasitism. This adds critical nuance to our understanding of the diverse mechanisms linking density with infection.

As hypothesized, we found strong negative relationships between NDVI (as a measure of resource availability) and parasite count – most likely because deer with access to more resources are more resistant to infection. Because immunity is costly, resource availability improves immune function in various taxa (González-Santoyo & Córdoba-Aguilar 2012; Budischak & Cressler 2018; Budischak *et al*. 2018; Hasik *et al*. 2021); therefore, the resource competition associated with higher density likely undermines immune resistance in the deer, leading to higher parasite counts. These findings demonstrate fundamentally that natural variation in density and its environmental effects can drive infection via susceptibility as well as via exposure, with effects that are likely multiplicative. Incorporating this variation into epidemiological models may help to accurately model the transmission and maintenance of pathogens in this and other systems. These findings are complementary to observations arising from resource supplementation in wild animal populations: generally, providing more resources drives greater contact rates, which drives increasing infection – counteracted by the improved resistance afforded by the resources and resulting immune function (Becker & Hall 2014; Budischak & Cressler 2018; Budischak *et al*. 2018; Hasik *et al*. 2021). This study therefore confirms that these interactions among density, resources, immunity, and infection occur in the wild as well as being dependent on human activities.

Supporting the argument that resistance is important in driving our effects, calves are the age class most heavily-infected with strongyles, likely due to their lack of an acquired immune response (Wilson *et al*. 2004; Ashby & Bruns 2018). Specifically, prior work has shown that not only are strongyle infections greater in calves (Albery *et al*. 2018), but also that immune defenses play a key role in defending from strongyle infection (Albery *et al*. 2020b). This lack of resistance may explain the strong density-dependence found for both strongyles and *E. cervi* in juveniles, as juveniles are unable to compensate for the increased exposure to parasites that comes with the increased density in the north of the study area. This observation agrees with our recent findings of greater density-dependence in young Soay sheep than in adults (Albery *et al*. 2024), indicating that density-dependent infections may generally be a larger problem for young (or weakly-immune) individuals across species.

Otherwise, the strong positive direct effects of density on infection status were likely indicative of greater levels of exposure, following the expectations of density-dependence theory (Anderson & May 1979; Arneberg *et al*. 1998; Detwiler & Minchella 2009). This effect was particularly expected for strongyles due to their direct life cycle: as deer density increases, more individuals shed larvae onto the pasture, creating higher larval abundances that then drive higher exposure rate. This represents useful observational evidence that within-population variation in density is associated with higher parasite counts. However, *F. hepatica* and *E. cervi*’s relationships may require further explanation, as both rely on an intermediate snail host to complete their life cycle. These results could be explained by the parasites’ life histories, as well as characteristics of both the deer and the study area.

The negative relationships between *F. hepatica* count and density and resource availability likely emerge from differences in habitat preference between deer and intermediate snail hosts. In the study area on Rum, the snails thought to act as intermediate hosts of *F. hepatica* are found in the stream alongside the track that runs along the western side of the study area, and the deer are less dense in this area compared to the north shore because the grazing in this area is generally lower quality (Fig. 1). This is the exact location where we found hotspots of *F. hepatica* infection (Fig. 1), such that intermediate host distributions may explain variation in *F. hepatica* infection because of shared causality (deer prefer to avoid marshes, while these aquatic snails prefer to inhabit them). This finding therefore serves to illustrate that density-dependence of infection can be highly confounded by environmental drivers and habitat selection; this is likely to be particularly important in environmentally-latent parasites and those with intermediate hosts.

Meanwhile, *E. cervi* relies on a terrestrial intermediate snail or slug host. On Rum we believe the relevant host is the heath snail (*Helicella itala*), which only occurs in the costal dunes in the north of the study area. This is not only where the greatest deer density occurs (Fig. 1), but also where *E. cervi* counts are highest. Mirroring *F. hepatica*, it could be that this is simply the result of both the deer and *H. itala* preferring this habitat. Another (non-exclusive) explanation is that this may be because the greater density of deer drives greater prevalence or intensity of infection in the snails, which could then increase the burden of *E. cervi* in the deer by increasing the rate at which they ingest *E. cervi* larvae. More data on the occurrence and distribution of the intermediate hosts of *F. hepatica* and *E. cervi* may better explain the spatial distribution of infection in the deer in the study area, as the incorporation of occurrence data of intermediate hosts can offer improved understanding of infection in the final host (Schols *et al*. 2021). This would require multiplying together spatial distributions of the host, the snails, and possibly environmental variables determining parasite development and transmission. Thus, an important next step for understanding parasitism in this system is to sample both the distribution and infection status of the aquatic and terrestrial snail hosts of *F. hepatica* and *E. cervi*, respectively.

Despite using a large amount of data from an exceptionally well-characterized system, our study had several important caveats. First and foremost, all of these data are observational, as it is impractical to interfere with the natural lives of the deer. We therefore cannot necessarily identify causation in the relationships that we have identified between the various population and environmental factors and the three parasites. Another consideration is that the analyses in this study are all based on fecal propagule counts, which are an emergent property of several separate processes. Namely, deer must first become exposed to infective larvae, those larvae must successfully establish within the host, and they must then reproduce. Internal processes such as immune defenses or variation in reproductive success among the parasites, especially among strongyle species, may obscure the true infection values. However, exposure to infective stages and resource-intensive immune resistance are both expected to be important contributors to observed fecal egg counts (McKenna 1981; Budischak *et al*. 2015; Watt *et al*. 2016), and our findings were broadly highly-supportive of the *a priori* hypotheses we set out. As such, we believe that the causal relationships we suggest to link density with infection via exposure and susceptibility are parsimonious and highly likely.

In conclusion, our study has revealed several important patterns relating the infection patterns of a long-lived mammal to the local environment and characteristics of the host population. These results not only support prior work in other systems (Gehman *et al*. 2017; Bolnick *et al*. 2020; Hasik & Siepielski 2022b), but highlight how important it is to compare relationships with the local environment to host metrics like population density when investigating infection. Further, by identifying the fine-scale spatial patterns of infection by three common helminth parasites, this study has provided an important first step toward the goal of understanding how parasites regulate host populations. Future studies could expand on this study by linking this variation to host fitness and survival, revealing how parasites themselves ultimately determine the distributions of host populations in space and time.

## Acknowledgements

We thank NatureScot for permission to work on the Isle of Rum and the many volunteers and researchers who have helped at the field site during this study. The field project is currently supported by the UK Natural Environment Research Council and the European Research Council. AZH benefit from the musical inspiration of Blood Command. This work was funded by a Leverhulme Research Grant (RPG 2022-220) awarded to JMP and GFA. GFA acknowledges funding from NSF DEB-2211287 and WAI (CBR00730).

---

**Figure S1.**
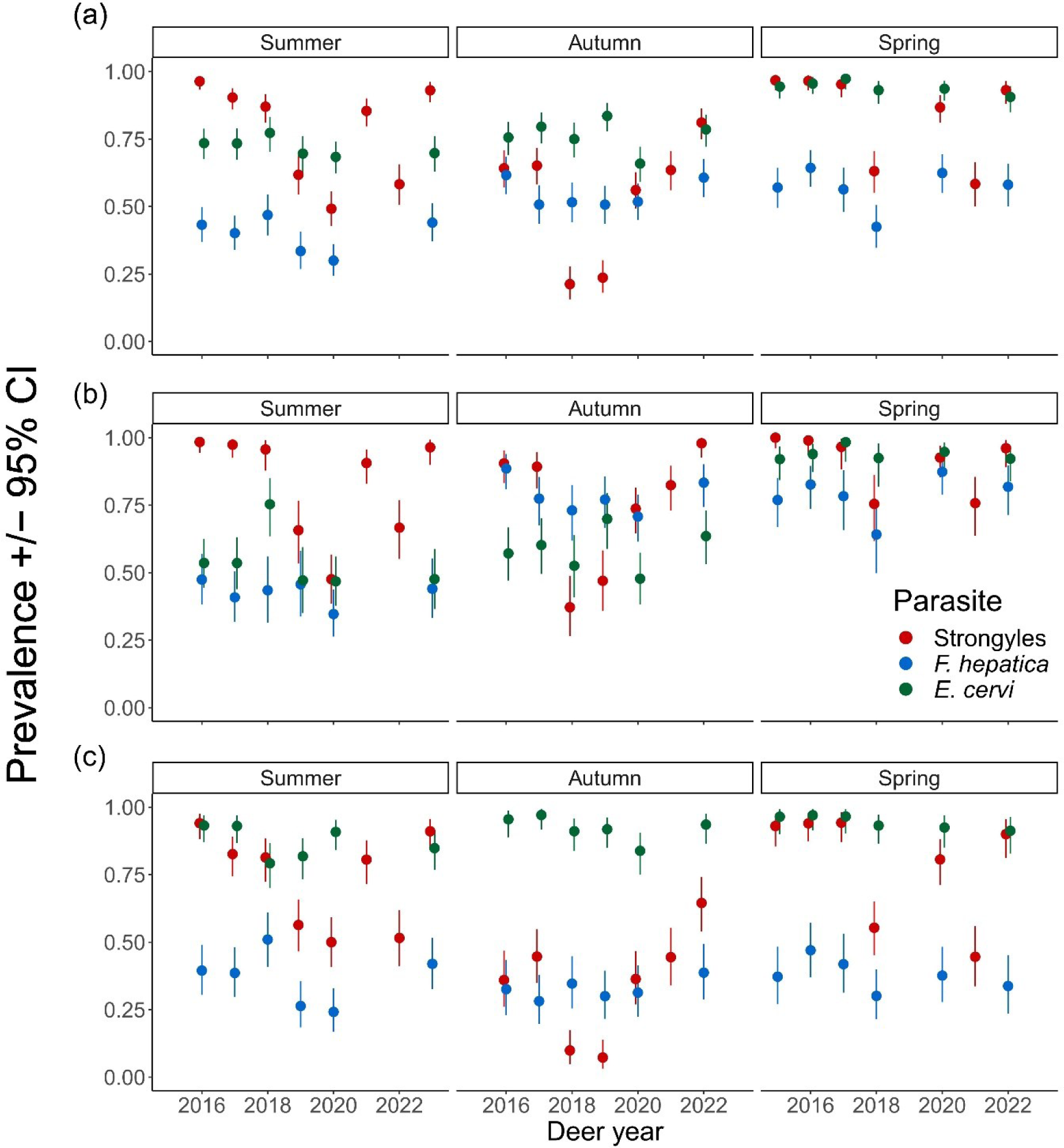
Plots of the seasonal prevalence patterns for each parasite for (a) all deer, (b) juveniles only, and (c) adult females only, with Deer year on the x-axis. The deer year runs from May 1^st^ to April 30^th^, so deer year 2016 ran from May 1^st^ 2016, with first sampling session in summer 2016, to April 30^th^ 2017. Points represent mean prevalence values, error bars denote 95% confidence intervals, and color denotes the parasite.

**Figure S2.**
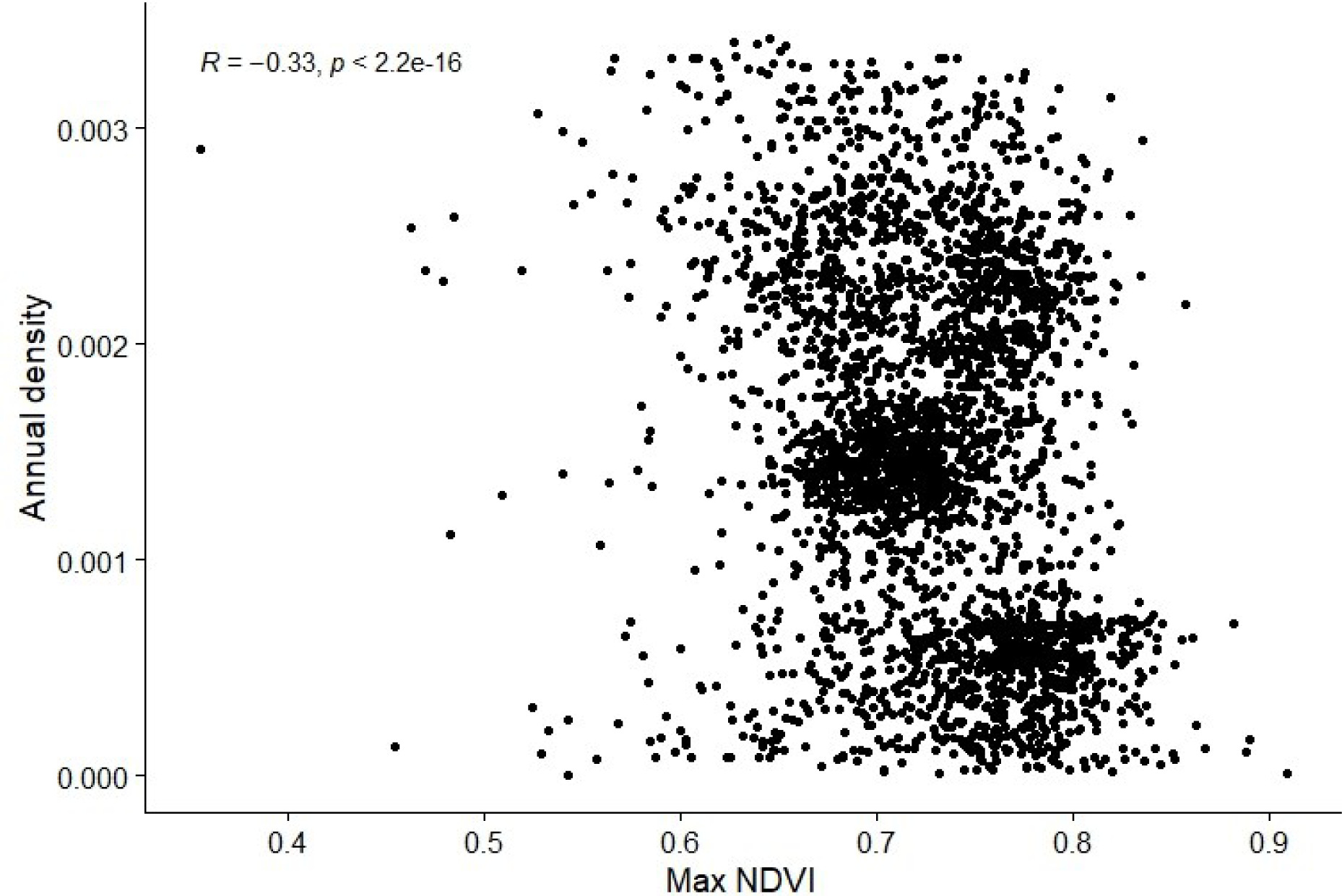
Plot of the correlation between annual max NDVI and mean annual density, with points representing individual deer.

**Figure S3.**
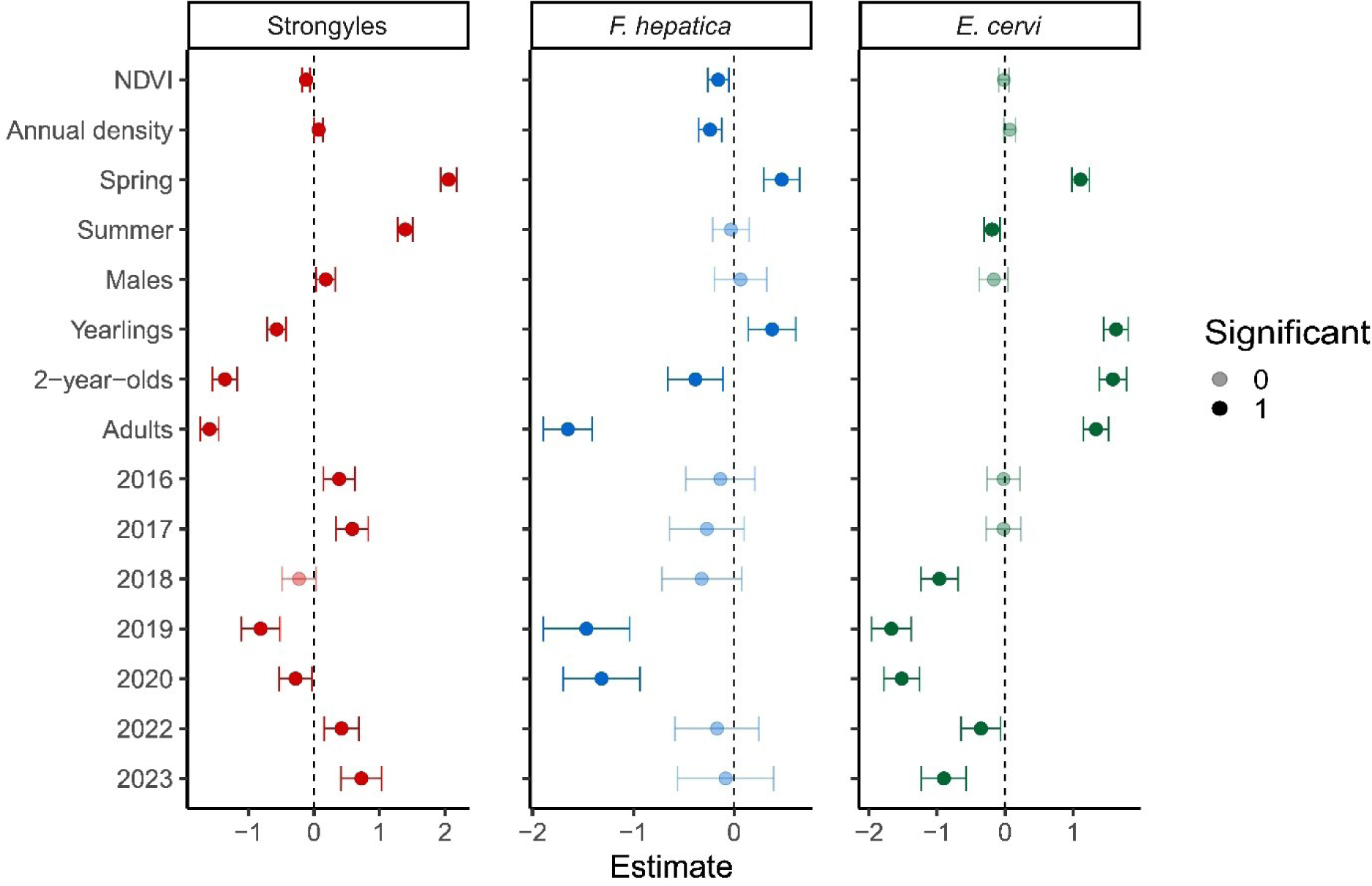
Forest plot representing the full model of the relationships between annual density, NDVI, season, age category, sex, and parasite counts for the dataset containing all deer, with panels for each parasite taxa. Points represent posterior estimates for mean effect sizes, error bars denote 95% credible intervals in standard deviations, and color denotes the parasite taxa. Significance of the effect size is denoted by the shading of the points.

**Figure S4.**
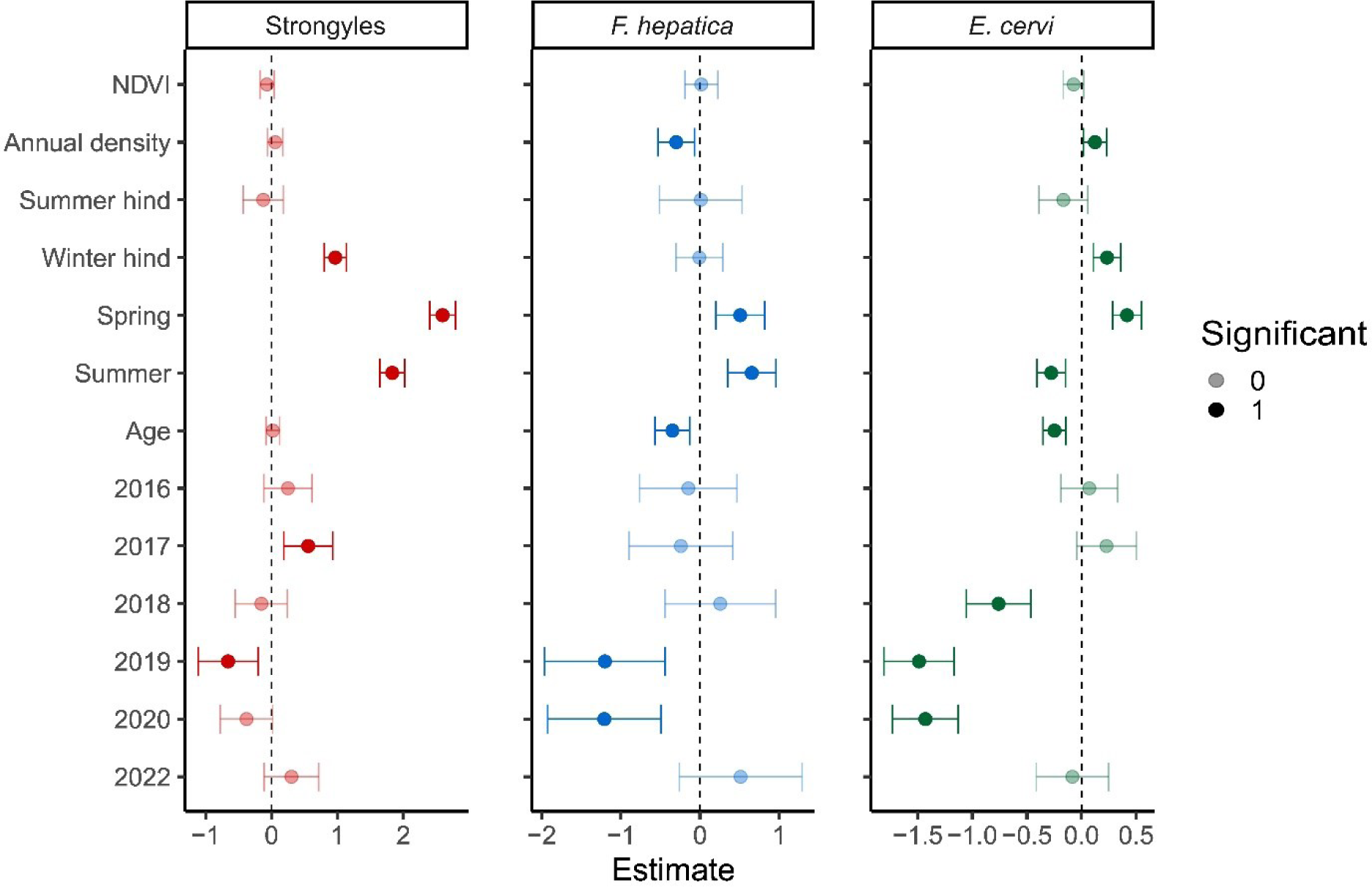
Forest plot representing the full model of the relationships between annual density, NDVI, reproductive status, age, season, and parasite counts for the dataset containing adult female only, with panels for each parasite taxa. Points represent posterior estimates for mean effect sizes, error bars denote 95% credible intervals in standard deviations, and color denotes the parasite taxa. Significance of the effect size is denoted by the shading of the points.

**Figure S5.**
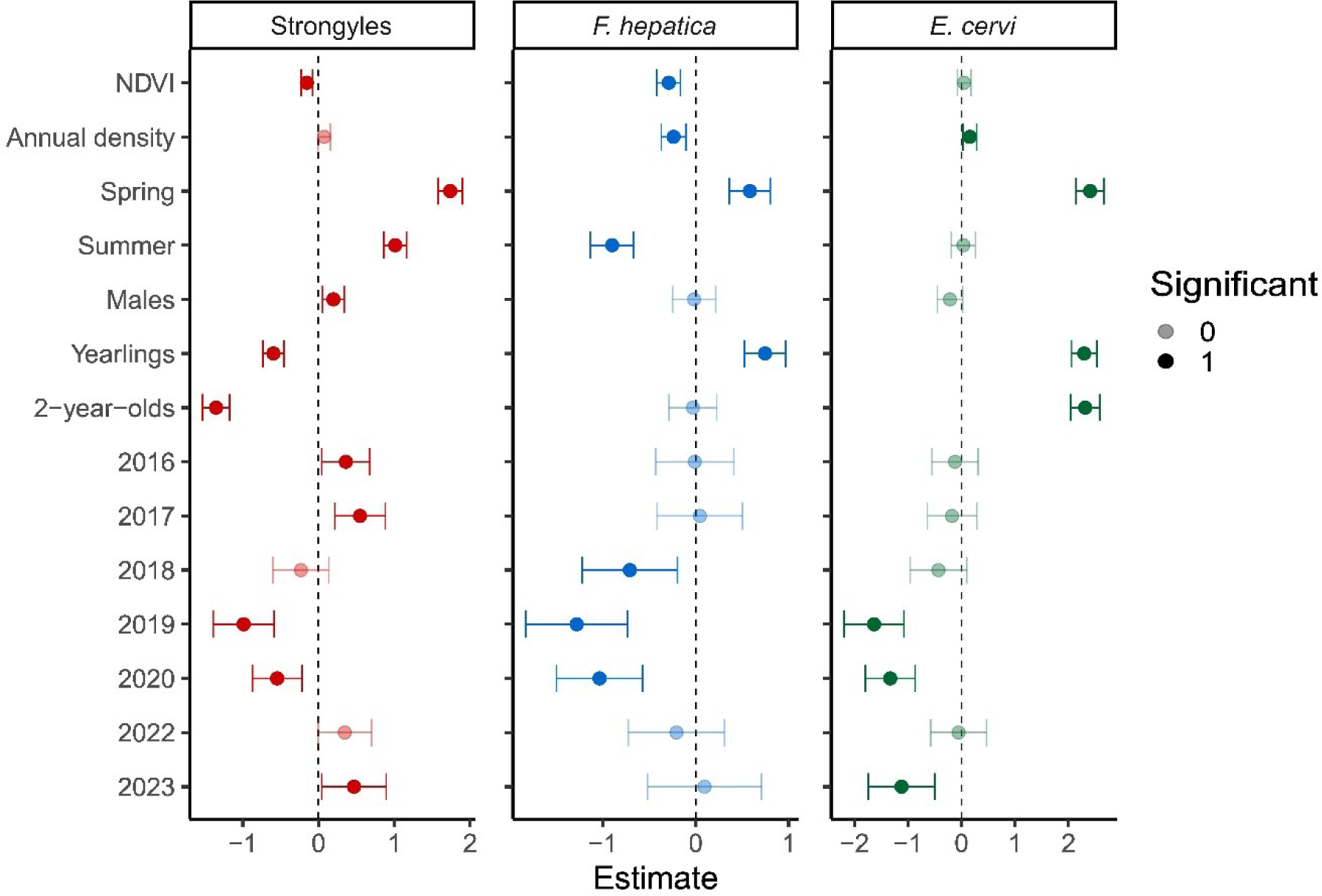
Forest plot representing the full model of the relationships between annual density, NDVI, season, age category, sex, and parasite counts for the dataset containing juveniles only, with panels for each parasite taxa. Points represent posterior estimates for mean effect sizes, error bars denote 95% credible intervals in standard deviations, and color denotes the parasite taxa. Significance of the effect size is denoted by the shading of the points.

